# *AUXIN RESPONSE FACTOR 6* (*ARF6)* and *ARF8* promote Gibberellin-mediated hypocotyl xylem expansion and cambium homeostasis

**DOI:** 10.1101/2020.11.13.381558

**Authors:** Mehdi Ben-Targem, Dagmar Ripper, Martin Bayer, Laura Ragni

## Abstract

During secondary growth, the thickening of plant organs, wood (xylem) and bast (phloem) are continuously produced by the vascular cambium. In Arabidopsis hypocotyl and root, we can distinguish two phases of secondary growth based on cell morphology and production rate. The first phase, in which xylem and phloem are equally produced, precedes the xylem expansion phase in which xylem formation is enhanced and xylem fibers differentiate. It is known that Gibberellins (GA) trigger this developmental transition via the degradation of DELLA proteins and that the cambium master regulator BREVIPEDICELLUS/KNAT1 (BP/KNAT1) and the receptor like kinases ERECTA and ERL1 regulate this process downstream of GA. However, our understandings on the regulatory network underlying GA-mediated secondary growth, are still limited.

Here, we demonstrate that DELLA-mediated xylem expansion is mainly achieved through RGA and GAI and that RGA and GAI promote cambium senescence. We further show that AUXIN RESPONSE FACTOR (ARF6) and ARF8, which physically interact with DELLAs, specifically repress phloem proliferation and induce cambium senescence during the xylem expansion phase. Moreover, the inactivation of *BP* in *arf6 arf8* background revealed an essential role for ARF6 and ARF8 in cambium establishment and maintenance. Overall, our results shed light on a pivotal hormone cross-talk between GA and auxin in the context of plant secondary growth.

## Introduction

Secondary growth, the increase in girth of plant organs, largely contributed to the success of land plants. Secondary growth is mainly driven by the vascular cambium, a post-embryonic meristem, which divides in a strictly bifacial manner, producing xylem (wood) inward and phloem outward (Ragni & Greb, 2018). In perennial dicotyledons, xylem tissue represents the principal form of biomass accumulation (Demura & Ye, 2010; Spicer & Groover, 2010). In addition, the vasculature contributes to the transport of assimilate, ions, water and signaling molecules and confers mechanical strength. Several studies suggest that the Arabidopsis hypocotyl is a valid model to study secondary growth and wood production (Chaffey *et al.*, 2002; Ragni & Hardtke, 2014), as it has been shown that key regulators are conserved between herbaceous and woody plants (Barra-Jimenez & Ragni, 2017) and extensive amounts of secondary xylem fibers and vessel elements are produced in the hypocotyl, with structural and ultra-structural characteristics similar to those of trees (Chaffey *et al.*, 2002).

In the Arabidopsis hypocotyl, secondary growth is characterized by two distinct growth phases. Upon termination of elongation at around 2-3 day-after germination, the hypocotyl grows only radially, xylem and phloem are produced at the same rate and the xylem comprises only vessels (first phase). In the second growth phase or the so called “xylem expansion” phase, xylem differentiates at higher rate than phloem and xylary fibers are formed (Sibout *et al.*, 2008). This developmental transition in the hypocotyl is induced by flowering in many herbaceous annual plants with rosette habitus including Arabidopsis (Ragni *et al.*, 2011). However, neither bolting nor flower specification per se are necessary. (Ragni *et al.*, 2011). Grafting experiments indicated the presence of a mobile signal, which at flowering is translocated from the shoot to the hypocotyl to induce xylem expansion (Sibout *et al.*, 2008). The phytohormone Gibberellin (GA) acts as the mobile cue that is transported at flowering to the hypocotyl where it drives DELLA degradation and promotes the xylem expansion program (Ragni *et al.*, 2011). Indeed, xylem expansion is enhanced in *della* quadruple mutants, in which GA signaling is constitutively turned on and abolished by the local expression of a non-degradable version of DELLA (Ragni *et al.*, 2011).

DELLA proteins are conserved components of the GA signaling pathway acting immediately downstream of the GA receptor, regulating a plethora of developmental processes and responses to abiotic stresses (Locascio *et al.*, 2013; Colebrook *et al.*, 2014; Davière & Achard, 2016). They control gene expression by sequestering transcription factors and repressors or by participating in transcriptional complexes (Locascio *et al.*, 2013). Consistently, they have been shown to interact with several transcription factor families (Marin-de la Rosa *et al.*, 2014) including: MYBs, NACs, and the AUXIN RESPONSIVE FACTORs/ARFs. For instance, RGA sequesters ARFs activators (ARF6, ARF7 and ARF8), but not the repressor ARF1 to regulate hypocotyl elongation (Oh *et al.*, 2014). Recently, it has been shown that RGA binds also to the transcription factor BREVIPEDICELLUS/KNAT1 (BP/KNAT1), a known secondary growth regulator (Liebsch *et al.*, 2014; Woerlen *et al.*, 2017; Felipo-Benavent *et al.*, 2018). Inactivation of BP leads to impaired cambial activity and absence of fiber differentiation. In addition, *bp* mutant do not form hypocotyl fibers even upon GA application indicating that BP acts also during the xylem expansion phase (Ikematsu *et al.*, 2017). Additionally ERECTA (ER) and ERECTA-LIKE1 (ERL1), two receptor like kinases, which are involved in many developmental processes such as stomata patterning and stem elongation (Shpak, 2013) regulate the transition to the xylem expansion in a BP-dependent manner (Ikematsu *et al.*, 2017).

The extent of xylem expansion varies among different natural accessions. In a pioneer study Sibout *et al.* identified several natural accessions with contrasting phenotypes (Sibout *et al.*, 2008), whereas we reported that the common laboratory strains Columbia (Col) and Landsberg *erecta* (Ler) greatly differ in the magnitude of xylem expansion. Ler has enhanced xylem expansion compared to Col, while Col displayed larger overall secondary growth than Ler (Ragni *et al.*, 2011; Sankar *et al.*, 2014). Surprisingly, the difference in xylem expansion among the two accessions cannot be explained by the inactivation of *ERECTA,* which occurs in the Ler ecotype (Ikematsu *et al.*, 2017). In a recent study, genome-wide association studies (GWAS) unraveled the role of another receptor like kinase SUPPRESSOR OF BRI1 /EVERSHED (SOBIR/EVR) during secondary growth (Milhinhos *et al.*, 2019). SOBIR, which acts together with ERECTA, prevents precocious fiber formation. Moreover, *SOBIR* expression is negatively regulated by BP (Milhinhos *et al.*, 2019). Nevertheless the GA signaling network underlying secondary growth is still poorly understood and so far BP/KNAT1 is the only known protein physically interacting with DELLA (Felipo-Benavent *et al.*, 2018), which regulates xylem expansion (Liebsch *et al.*, 2014; Woerlen *et al.*, 2017; Felipo-Benavent *et al.*, 2018).

In this work, we show that RGA and GAI are the main members of the DELLA family regulating the transition to the xylem expansion phase, highlighting their role in modulating cambium senescence (decrease of cambium activity at the last plant developmental phase). We expanded the GA downstream network by integrating ARF6 and ARF8 as two DELLA-interacting proteins and we demonstrated that ARF6 and ARF8 repress phloem proliferation and regulate cambium activity. Interestingly, *arf6 arf8* double mutants show ectopic phloem proliferation only after the onset of xylem expansion phase and delayed cambium senescence, highlighting their specific function in this developmental transition. In addition, the inactivation of *BP* in *arf6 af8* background results in a nearly total arrest of secondary growth, unmasking a possible ARF6 and ARF8 function in establishing and maintaining active vascular cambium.

## Experimental procedure

### Plant material and growth

The majority of the lines used are in Col background unless otherwise specified in the text/figure. All plants used for confocal microscopy are in Col background, grown in vitro (1/2 MS with 0.1% of sugar) in continuous light. For all the other experiments, plants were grown in soil in long day conditions (16 hours light versus 8 hours dark) and the sampling time is stated in the text/figure. *nph4-1, arf6-2, arf8-3, arf6-2 nph4-1 arf8-3/+, pRPS5a::GAL4* and *pUAS::MIR167* and the F1 issued from the cross between them were kindly provided by Jason Reed (University of North Carolina, USA) and described in (Nagpal *et al.*, 2005) (Stowe-Evans *et al.*, 1998). *arf8-7* (Gutierrez *et al.*, 2009) *ARF6:GUS* and *ARF8:GUS* (Nagpal *et al.*, 2005; Wu *et al.*, 2006) were kindly provided by Catherine Bellini (Umeå university, Sweden). The double *rga-24 gai-t6* was a gift from Markus Schmid (Umeå university, Sweden). The *dellako, er-105* and *SUC2:GUS* were previously described in (Feng *et al.*, 2008) (Torii *et al.*, 1996) and (Schulze *et al.*, 2003). *rga-28, gai-td1* and the double *rga-28 gai-td1* (Plackett *et al.*, 2014) were provided by Stephen Thomas (Rothamsed Research). *bp-9* (Pautot *et al.*, 2001), *BP:GUS* (Pautot *et al.*, 2001) are gift from V.Pautot (INRA Versailles), *RGA:rgaD-GR, GAI:gaiD-GR, rga-24, gai-t6* were previously described in (Ragni *et al.*, 2011) *ARF6::NLS-3xGFP* (N67078), *ARF7::NLS-3xGFP* (N67080), *ARF8::NLS-3xGFP* (N67082) *arf6-1*(N24606) *and arf8-2*(N24608) (Okushima *et al.*, 2005) were ordered from NASC. *RGA:rgaD-GR, GAI:gaiD-GR* in Col-0 background were obtained by backcrossing 6 times the original lines to Col-0. *arf6-1 arf8-2/+, bp-9 arf6-2 arf8-3, BP:GUS arf6-2 arf8-3, arf6-2 rga-28 gai-td1/+ arf8-3* were generated by crossing and genotyping. Primers for genotyping are listed in Table S1.

### GA and Dexamethasone Treatment

Induction of soil grown *RGA:rgaD-GR,* or *GAI:gaiD-GR* plants in Ler or Col-0 background was achieved by watering with a 10 μM dexamethasone solution three times per week until hypocotyl sampling. Plants grown invitro, were treated by submerging in a liquid 10 μM dexamethasone (dex) ½ Murashige and Skoog solution for 3 hours upon flowering. For GA and dex on soil treatments, plants were watered from flowering on, with a 100 μM GA3/ 10 μM dex or mock solution at similar frequency three times per week until hypocotyl sampling according to the experiment.

### Molecular Cloning

The RGA promoter was amplified with the primers: A-pRGA F (AACAGGTCTCAACCTTATAACCTCATCCATCTATAG) and Br-pRGA R (AACAGGTCTCATGTTTCAGTACGCCGCCGTCGAGAG). The BsaI site inside the pRGA promoter was removed by overlapping PCRs and cloned in pGG-A0. The GAI promoter was amplified with the primers: A-pGAI F (AACAGGTCTCAACCTTGGGACCACAGTCTAAATGGCGT) and Br-pGAI R (AACAGGTCTCATGTTGGTTGGTTTTTTTTCAGAGATGGA) and cloned in pGG-A0. The promoters were assembled in pZ03 with the available published modules to obtain RGA:NLS-GFP-GUS and GAI: NLS-GFP-GUS by Green Gate technologies (Lampropoulos *et al.*, 2013).

### Histology and stainings

Thin plastic cross-sections were obtained from plastic embedded material using TECHNOVIT 7100 (Heraeus Kulzer) and a Leica RT microtome as described in (Barbier de Reuille & Ragni, 2017). GUS essay was performed according to the protocol described in (Wunderling *et al.*, 2018). Vibratome sections (50–80 μm) were obtained via a Leica VT-1000 vibratome, from hypocotyls embedded in 6% agarose block, slices were collected in water in microscope slides, stained and/or imaged. For Phloroglucinol staining, a ready solution (VWR, 26337.180) was applied directly to the section. Images were taken with a Zeiss Axio M2 imager microscope. For sections of fluorescent lines the ClearSee protocol described in (Ursache *et al.*, 2018) was applied. Briefly, the hypocotyls were first fixed in a 4% PFA solution with 0.01% Triton for 1 hour. and then embedded in 5% Agarose for vibratome cutting. Vibratome sections are then directly collected in 1x PBS solution. After incubation, the 1x PBS solution is removed and replaced by ClearSee solution. The sections are subsequently kept at room temperature in ClearSee solution for at least 1 to 2 days. Finally, the samples are incubated for 20 min at RT in 0,05 % Calcofluor White then washed and mounted on slides in ClearSee solution (Kurihara *et al.*, 2015).

### Images acquisition

A Zeiss Axio M2 imager microscope or a Zeiss Axiophot microscope was used to take images at different magnifications of vibratome sections (50–80 μm) as well as 5 μm sections stained with 0.1% toluidine blue solution. Pictures of vibratome sections of fluorescent lines were acquired using the Zeiss LSM880 confocal microscope. GFP ex. 488 em. 490-520. Calcofluor White ex 404 em: 430-450.

### Image analyses and statistical analyses

The total hypocotyl cross section area, the xylem area and the fiber area were analyzed using ImageJ software as previously described (Sibout *et al.*, 2008; Wunderling *et al.*, 2017). Statistical analyses were performed using IBM SPSS Statistics version 24-25 (IBM). We first tested all datasets for homogeneity of variances using Levene’s Test of Equality of Variances. For multiple comparison, we calculated the significant differences between each dataset using One way ANOVA with Tamhane’s post hoc (equal variance not assumed) or a Bonferroni correction (equal variance assumed). The significance threshold was set to p-value < 0.05. For comparing two groups, we used Welch’s t-test (not homogenous variance) or a Student’s t-test (homogeneous variance), P values are indicated in the figure legends.

## Results

### GA signaling controls hypocotyl secondary growth, mostly through RGA and GAI, in Ler and Col-0 ecotypes

We previously showed that *della* quadruple mutants are characterized by an increased xylem expansion and fiber production (Ragni *et al.*, 2011), however, the specific contribution of each DELLA during secondary growth, is unknown. Thus, we analyzed *della* single knock-out mutants. Only *rga* mutants exhibited a mild but significant increase of xylem to total area ratio (from now on referred as xylem occupancy) in comparison to wildtype (Figure S1), suggesting a certain degree of redundancy. As *GAI* has been shown to play overlapping roles with *RGA* in several developmental processes (Dill & Sun, 2001; Cheng *et al.*, 2004), we investigated the *rga-24 gai-t6* double mutants. *rga-24 gai-t6* mutants showed a strong increase in xylem occupancy and fiber formation (Figure S1c-d), whereas knocking out all *DELLAs (dellako)* only slightly enhanced the *rga gai* phenotype, suggesting that *RGA* and *GAI* are the major regulators of xylem expansion (Figures S1c-d).

Plakett el al. showed that *rga gai* double mutants are sterile in Col background, whereas they set seeds in Ler background (Plackett *et al.*, 2014). As Ler and Col also greatly differ in secondary growth morphodynamics with Col displaying more overall secondary growth and Ler more xylem expansion (Ragni *et al.*, 2011; Sankar *et al.*, 2014), we investigated the vascular phenotype of *rga gai* in both ecotypes. Both *rga-24 gai-t6* (Ler background) and *rga-28 gai-td1* (Col background) displayed enhanced xylem expansion, compared to their wild-type (WT) counterparts, however, the absolute values reflect the differences between the two ecotypes with *rga-24 gai-t6* having 50% xylem occupancy (in Ler) and *rga-28 gai-td1* only 30% (in Col) (Figures 1a-b). Interestingly, the triple mutant *rga-28 gai-td1 er-105* was undistinguishable from the *rga-28 gai-td1* double mutant in terms of xylem expansion (Figures 1a-b) pointing out that the difference between the two double mutants is not due to the inactivation of *ERECTA* in Ler background. Consistently, it was previously shown that, the loss of function of *ERECTA* (in Col background) does not increase xylem expansion (Ikematsu *et al.*, 2017) (Figures 1a-b). To corroborate our results, we studied the effect of the induction of a non-degradable version of RGA and GAI *(GAI:gaiD17-GR* and *RGA:rgaD17-GR),* in both backgrounds. To this extent, the original *GAI:gaiD17-GR* and *RGA:rgaD17-GR* Ler lines (Ragni *et al.*, 2011) have been backcrossed 6 times to Col background. Induction, upon flowering, of a DELLA version that cannot be degraded in the presence of GA resulted in reduced xylem occupancy and fiber formation in both accessions (Figures S2a-b). Altogether these results suggest that in Ler background a xylem promoting or phloem repressing factor acts in a DELLA- and ERECTA-independent pathway to promote the xylem expansion phase. Thus, to easily characterize the GA downstream pathway, we mainly exploited the Col background.

**Fig. 1.**
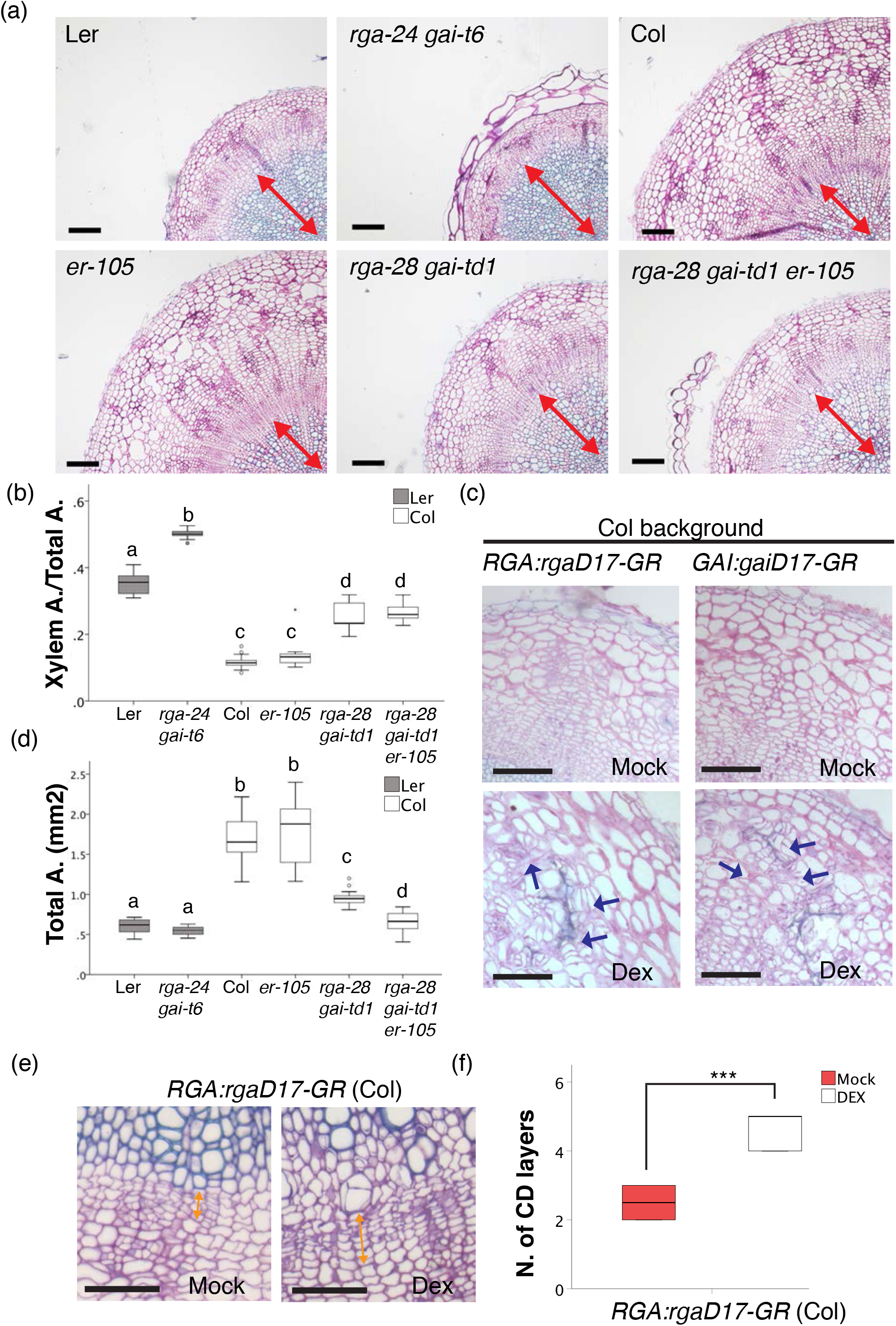
GA signalling controls hypocotyl secondary growth, mostly through *RGA* and *GAI,* in Ler and Col ecotypes. (a) Plastic cross-sections of 10 day-after-flowering (daf) hypocotyls showing xylem expansion in WT (Ler and Col background), *rga-24-gai-t6* (in Ler background), *rga-28 gai-td1* (in Col background), *er-105* (in Col background) and *rga-28 gai-td1 er-105* (in Col background). Double-headed red arrows indicate the xylem. Black scale bars:100μm (b) Quantification of Xylem Area/Total area of the experiment showed in (a). Letters in the graphs refer to individual groups in a one-way ANOVA analysis with a post-hoc multiple group T-test (n=11-19). (c) Plastic crosssections of 15daf hypocotyls of *rga:rgaD17-GR* and *gai:gaiD17-GR* in Col. Plants were treated with 10μM DEX from flowering until sample collection. Blue arrows indicate ectopic cell divisions in the phloem. Black scale bar: 100μm. (d) Quantification of Total Area of the experiment showed in (a). Letters in the graphs refer to individual groups in a one-way ANOVA analysis with a post-hoc multiple group T-test (n=11-19). (e) Magnification of (c) showing the cambium of *rga:rgaD17-GR* mock and treated hypocotyls. Double-headed orange arrows indicate the cambium. Black scale bar: 20μm. (f) Quantification of the number of cambium derivative layers (CD layers) of the experiment showed in (e). T-test (n=6, ***: p<0.001).

In Col ecotype, we observed further secondary growth phenotypes when we perturbed GA signaling possibly due to the absence of “the xylem promoting factor”. Induction of *gaiD17* or *rgaD17* upon flowering, promotes ectopic cell divisions in the phloem and resulted in an increase in total hypocotyl area (Figures 1c and S2c). Consistently, *rga-28 gai-td1* mutants (Col background) are characterized by a reduction in total hypocotyl area (Figure 1d). This led us to further investigate cambium activity in the context of xylem expansion. For this purpose, we measured the number of cambial derivative layers (CD layers), at the onset (5 daf), during early and late xylem expansion (15-30 daf). We observed that in WT hypocotyls during the late xylem expansion phase, the number of CD layers decrease, ultimately leading to cambium consumption at plant senescence, from now on this phenomenon will be referred to as cambium senescence (Figure S2d). By contrast, the number of cambium cell layers in dex-treated *RGA:rgaD-GR* plants was increased compared to mock (20 daf) indicating that DELLAs delay cambium senescence (Figures 1e-f). This is in line with the fact that *rga-28 gai-td1* double mutants displayed less cambial layers than WT (10 daf) (Figures S2e-f), indicating that DELLA proteins delay cambium consumption during senescence.

### BP is not the only factor required for fiber formation

BP has been shown to regulate fiber differentiation during xylem expansion through its physical interaction with DELLA (Felipo-Benavent *et al.*, 2018) *bp* mutants (in Col ecotype) lack fiber differentiation and are insensitive to GA (Ikematsu *et al.*, 2017). This is in contrast to what we observed in *bp* mutants in Ler ecotype (*bp-1*), which can still partially respond to GA application (Figures S3a-b). *bp-1* mutant plants were able to form fibers albeit at lower extent and only very late in development (20 daf), suggesting that fiber formation is possible in the absence of BP, in a background in which xylem expansion is enhanced (Figure S3a).

To better clarify this issue, we investigated whether the inactivation of RGA and GAI rescues the *bp* phenotype (in Col background). In *rga gai bp-9,* plant height phenotype was partially rescued (Figure S3c) and we observed fiber formation at the time of plant senescence, nevertheless, the altered xylem occupancy was not rescued by the inactivation of RGA and GAI. Altogether these results indicate that BP is not the only factor required for the xylem expansion phase (Figures S3d-f).

### ARF6, ARF7 and ARF8 expression patterns overlap with RGA and GAI during hypocotyl secondary growth

In the quest for GA downstream factors controlling hypocotyl secondary growth, we thought that ARF6, ARF7 and ARF8 are promising candidates, as they physically interact with RGA and GAI (Oh *et al.*, 2014) and auxin is a key regulator of vascular patterning and xylem differentiation (Ragni & Greb, 2018). To investigate whether ARF6, ARF7 and ARF8 regulate the xylem expansion phase through the interaction with DELLA, we first checked whether their expression patterns overlap with RGA and GAI during secondary growth. *RGA* and *GAI* are broadly expressed in the hypocotyl before flowering (Figure S4a), whereas at flowering time, *RGA* and *GAI* expression gets progressively restricted to the phloem poles. During xylem expansion, RGA and GAI are still expressed at the phloem poles albeit at lower levels (10 day-after-flowering). Consistently RGA protein accumulation was reported in the phloem (Felipo-Benavent *et al.*, 2018)(Figure 2a, b). Interestingly, *ARF6, ARF7* and *ARF8* expression was also detected in the phloem throughout secondary growth, *ARF6* and *ARF7* expression was broader and encompassed the cambium and differentiating xylem respectively, similar to their expression pattern in the root, described by (Smetana *et al.*, 2019) (Figures 2c-e). In addition, ARF6 and ARF7 and to a lesser extent ARF8 expression was maintained also after flowering. To conclude, *RGA, GAI, ARF6, ARF7* and *ARF8* expression patterns overlap in the phloem, therefore fulfilling the requirement that these factors could physically interact also during secondary growth.

**Fig. 2.**
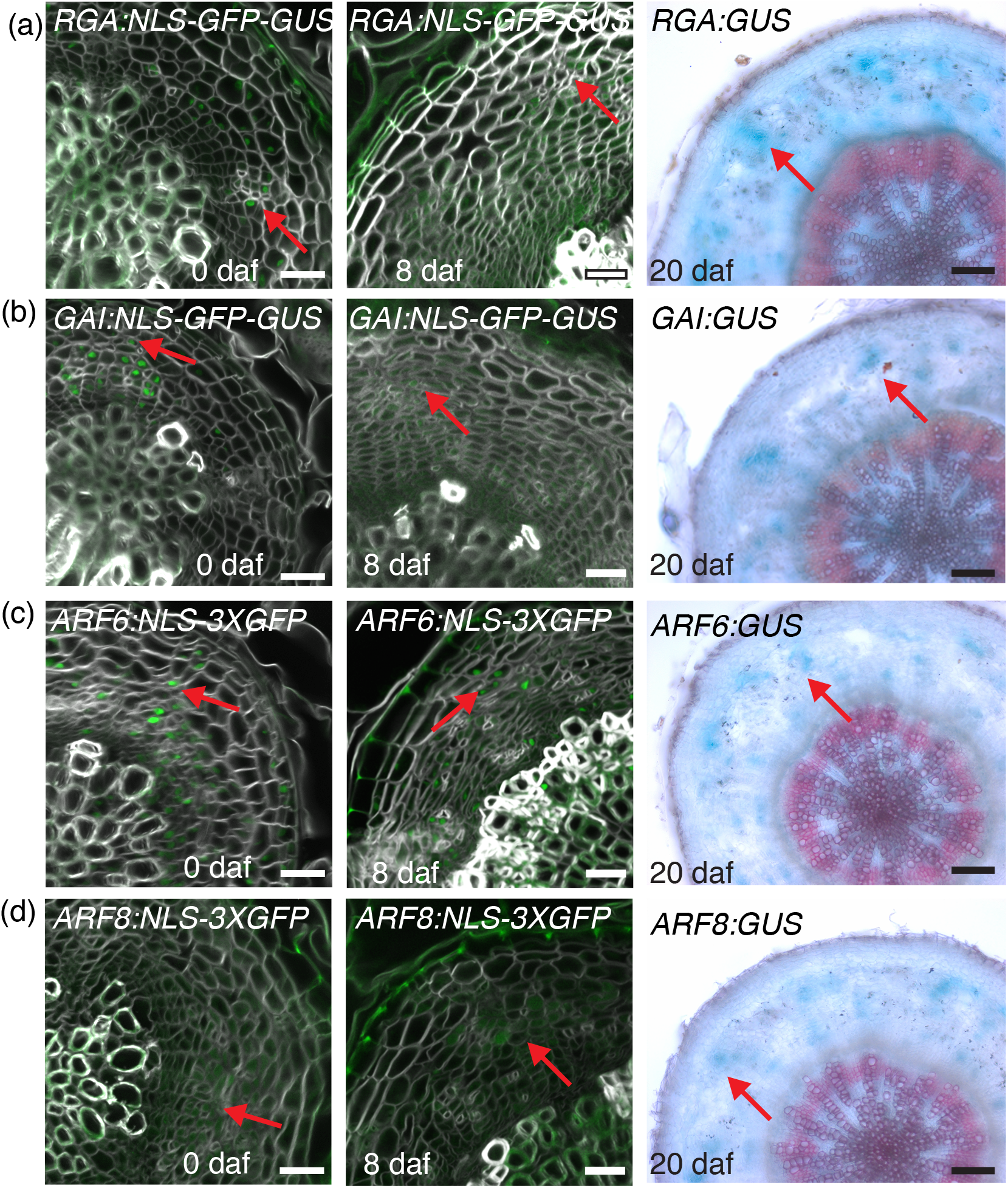
*ARF6* and *ARF8* expression pattern overlaps with *RGA* and *GAI* during hypocotyl secondary growth. (a-d) Hypocotyl vibratome cross-sections at 0, 8 and 20 day-after-flowering. Left and middle panels: confocal images of sections cleared with ClearSee and stained with Calcofluor-White showing GFP signal in the nuclei (red arrows). Right panels: GUS assay on vibratome sections stained with Phloroglucinol (red arrows indicates the phloem). (a) *RGA:NLS-GFP-GUS* (left and middle panels) and *RGA:GUS* (right panel). (b) *GAI:NLS-GFP-GUS* (left and middle panels)and *GAI:GUS* (right panel). (c) *ARF6:NLS-3xGFP* (left and middle panels) and *ARF6:GUS* (right panel). (d) *ARF8:NLS-3xGFP* (left and middle panels) and *ARF8:GUS(right* panel). (a-d) White scale bar: 20 μm and black scale bars: 20μm.

### ARF6, ARF8 and to a lesser extent ARF7 promote xylem expansion, repressing phloem proliferation

To further investigate the ARF-DELLA genetic interaction during secondary growth, we tested whether *arf* mutants displayed altered xylem expansion. *arf7*/*nph4* mutants are characterized by reduced total hypocotyl area but did not show any xylem expansion phenotype (Ragni *et al.*, 2011). *arf6 (arf6-1* and *arf6-2)* and *arf8 (arf8-2* and *arf8-3)* single mutants showed a slight but significant inhibition of xylem expansion and less fibers compared to the WT background (Figs. 3 a-d and S4c). Whereas the *arf6 arf8 (arf6-2 arf8-3* and *arf6-1 arf8-2)* double mutants displayed a dramatic decrease in xylem occupancy and absence of fiber accumulation until very late stages of plant growth (30-40 days after flowering in our growth conditions) (Figures 3a-d and S4c). Interestingly in *arf6 arf8* double mutants, we observed the presence of phloem fibers at plant senescence, which rarely occur in WT plants (Figure 3b), an increased number of phloem poles and ectopic cell divisions at phloem poles (Figure 3e and S4d) reminiscent of the phenotypic consequences of impaired GA signaling in Col background (Figure 1c). To confirm that this phloem phenotype is due to the inactivation of *ARF6* and *ARF8,* we took advantage of the fact that the expression of *ARF6* and *ARF8* is tightly regulated by miR167a (Wu *et al.*, 2006; Gutierrez *et al.*, 2009). Supporting the role of *ARF6* and *ARF8,* we observed ectopic cell divisions in the phloem of F1 plants, in which we ectopically expressed miR167a *(UAS::miR167a xRPS5a::GAL4)* (Figure S4e). These results suggest that the transition to the xylem expansion phase relies on reducing cambium cell differentiation into phloem, resulting in a shift towards xylem differentiation.

**Fig. 3.**
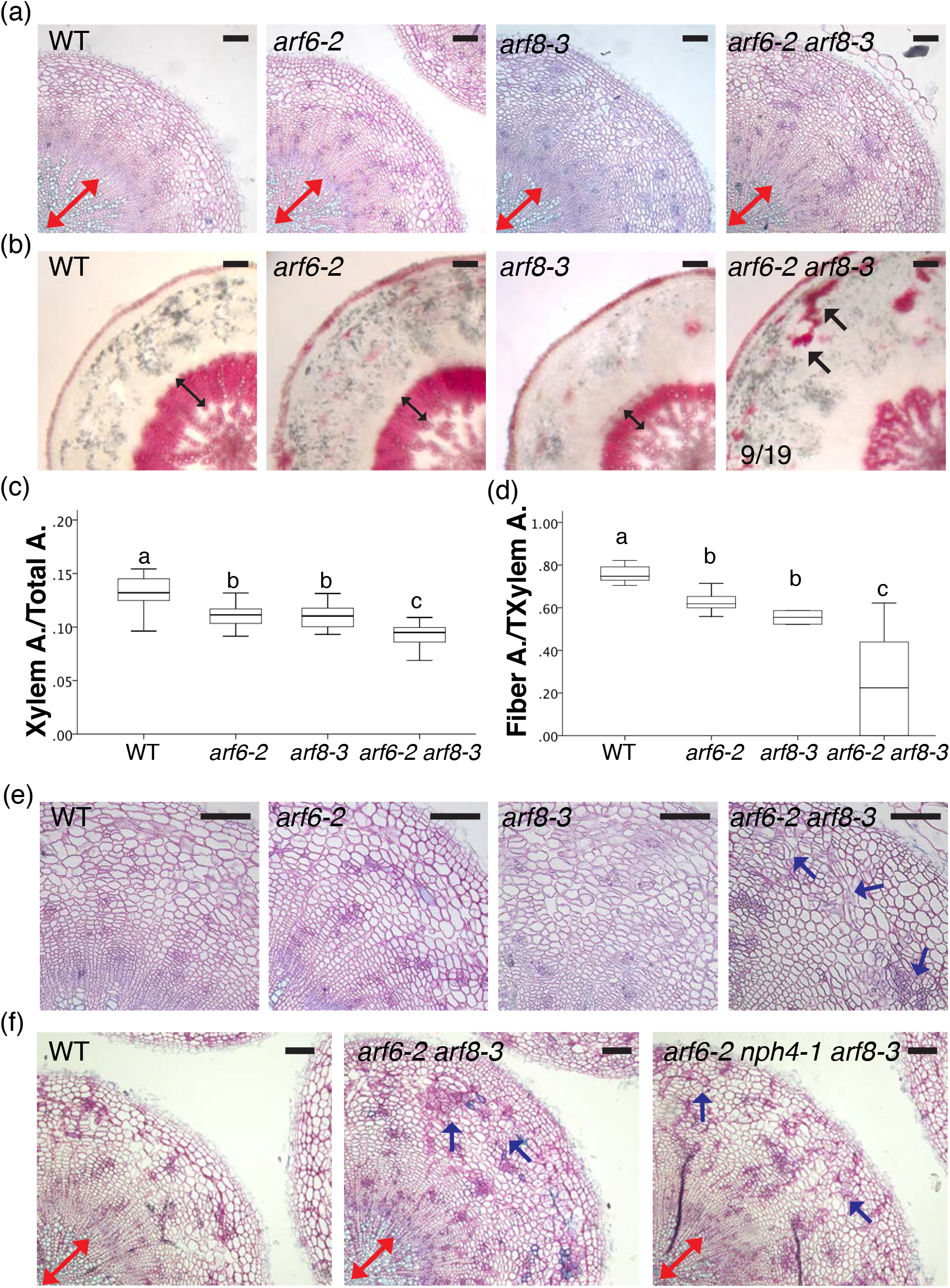
ARF6, ARF8 and to a lesser extent ARF7 promotes xylem expansion, repressing phloem proliferation. (a) Plastic hypocotyl cross-sections of 10 day-after-flowering (daf), showing reduced xylem expansion in *arf6*, *arf8* and *arf6 arf8* mutants compared to WT (Col). (b) Phloroglucinol stained vibratome sections of 40 daf hypocotyl, showing reduced fibers formation in *arf6*, *arf8* and *arf6 arf8* mutants compared to WT (Col). (c) Quantification of the Xylem Area /Total area ratio of the experiment showed in (a). Letters in the graphs refer to individual groups in a one-way ANOVA analysis with a post-hoc multiple group T-test (n=9-17). (d) Quantification of Fiber Area/Xylem Area ratio of the experiment showed in (b). Letters in the graphs refer to individual groups in a one-way ANOVA analysis with a post-hoc multiple group T-test (n=2-18). (e) Magnifications of (a) showing the phloem region. (f) Plastic hypocotyl cross-sections of 10 daf, showing reduced xylem expansion and ectopic cell divisions in the phloem in *arf6*, *arf8* double and *arf6 nph4/arf7 arf8* triple mutants compared to WT. Black scale bars: 100μm. Double-headed red arrows indicate the xylem, double-headed black arrows indicate xylem fibers, black arrow indicate phloem fibers and blue arrows indicate ectopic divisions in the phloem.

We next wondered about the temporal dynamics of ARF6 and ARF8 DELLA interaction: if RGA sequesters ARFs during the first phase of secondary growth, then we would expect that *arf6 arf8* mutants show strong changes in secondary growth only after flowering. Consistently with this hypothesis, *arf6 arf8* hypocotyls were undistinguishable from WT plants at flowering time *(arf6 arf8* double mutants flower at the same day as WT in long day conditions) (Figures S4f). After flowering (from ≡10 daf), however, xylem occupancy starts to increase in wild-type plants, whereas in *arf6 arf8* (at 15 daf) it was unchanged and only increased mildly at 30 daf (Figures S5a-b). By contrast, we remarked that total hypocotyl area was bigger in *arf6 arf8* double mutants compared to WT (Figures S5c). This could be explained by prolonged cambium activity, in *arf6 arf8* mutants. At the onset of xylem expansion and during early xylem expansion phase (5-15daf), the number of cambial derivative layers was similar in WT and *arf6 arf8* mutants (@ 7 layers), whereas in the late xylem expansion phase (at 30 daf), cambial activity was decreased in WT but not in *arf6 arf8* plants (≅ 4 layers vs ≅ 6 layers), indicating that *ARF6* and *ARF8* promote cambium consumption (Figures S5d, e).

Finally, we wondered whether the inactivation of ARF7 could enhance the *arf6 arf8* phenotype. *arf7/nph4 arf6 arf8* displayed a further reduction of xylem occupancy (Figure S5f), compared to the double mutant, however the ectopic phloem cell divisions phenotype was not enhanced (Figure 3f and S5g), suggesting only a minor contribution by *ARF7.*

### ARF6, ARF7 and ARF8 act mainly downstream of GA signaling to control xylem expansion

To further confirm the interaction between DELLA and ARFs and its significance in controlling GA mediated xylem expansion, we tested whether GA responses are attenuated in *arf6 arf8* double mutants. This was achieved by exogenous GA application and by inactivating *RGA* in *arf6 arf8* background. As previously shown, GA treatment enhances fiber production and increases xylem occupancy in WT (Ragni *et al.*, 2011) (Figures 4a-c), by contrast *arf6 arf8* mutants were only partially able to respond to GA as indicated by the mild increase in xylem occupancy and fiber production, (Figure 4a-c). Moreover, in *arf7/nph4 arf6 arf8* triple mutants, GA treatment, did not increase xylem occupancy and fiber accumulation was abolished in the majority of the plants (Figures 4a-c). In line with this observation, *rga ar6 arf8* triple mutants showed decreased xylem occupancy and no fiber production compared to *rga* single mutant (Figures S6a-c). Similarly, *arf6 arf8 rga gai/+* showed less xylem occupancy and fibers compared to *rga gai/+* (Figures S6a-c). Over all, these results confirm that *ARF6, ARF7* and *ARF8* are required to mediate GA dependent xylem expansion. The partial restoration of the *arf6 arf8* phenotype caused by DELLA inactivation or GA treatment could be explained by the additional role of ARF7 in xylem expansion or ectopic BP activity in *arf6 arf8* mutants (see below).

**Fig. 4.**
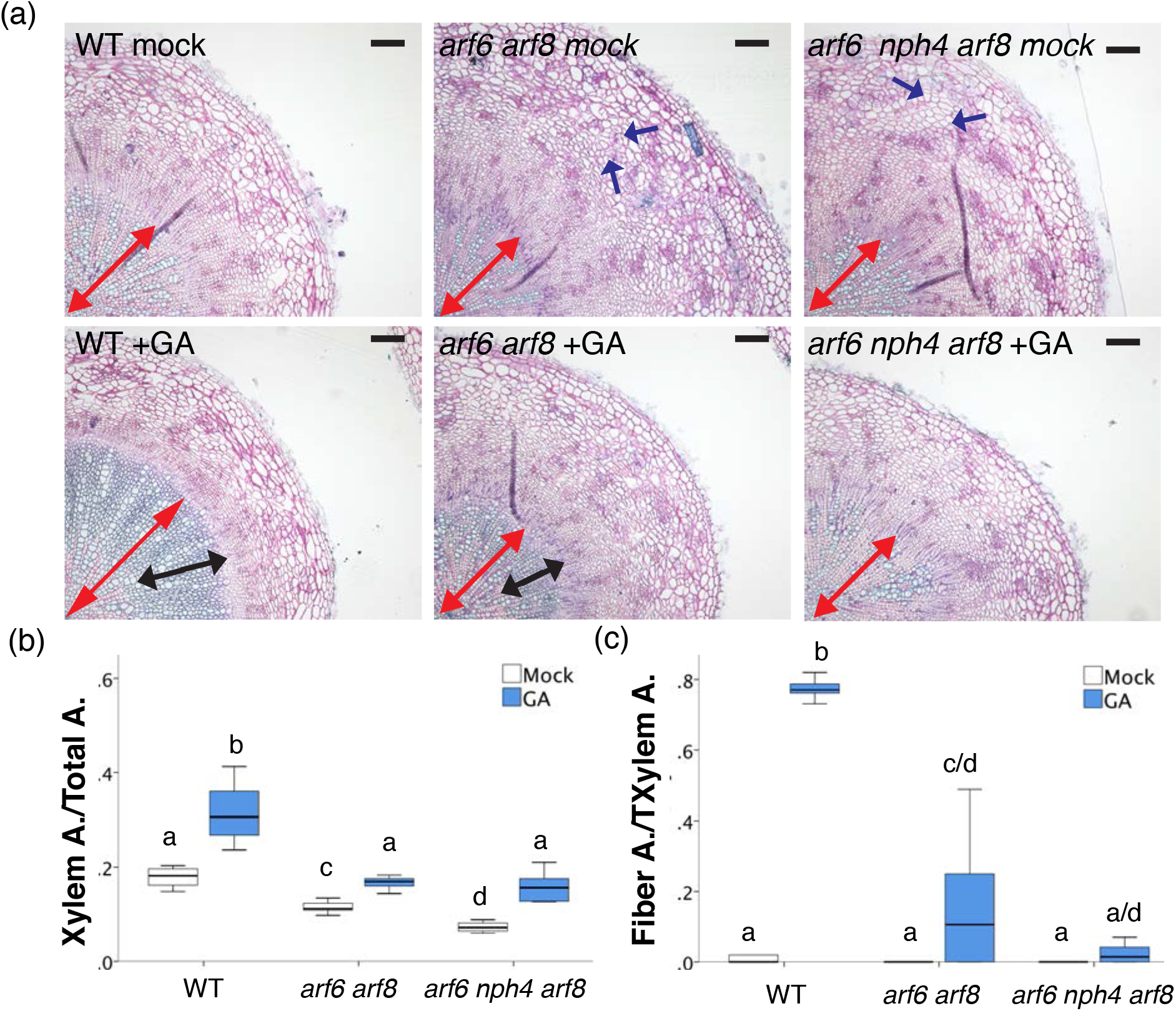
ARF6, ARF7 and ARF8 act mainly downstream GA signalling to control xylem expansion. (a) Plastic hypocotyl cross-sections of 10 day-after-flowering, showing xylem expansion in WT (Col), *arf6 arf8* and *arf6 nph4/arf7 arf8* plants treated with Mock or 10μM GA solution. (b) Quantification of Xylem Area/Total area ratio of the experiment showed in (a). Letters in the graphs refer to individual groups in a one-way ANOVA analysis with a post-hoc multiple group T-test n=4-13). (c) Quantification of Fiber Area/Xylem Area ratio of the experiment showed in (a). Letters in the graphs refer to individual groups in a one-way ANOVA analysis with a post-hoc multiple group T-test n=4-13). Black scale bars: 100μm. Double-headed red arrows indicate the xylem, double-headed black arrows indicate xylem fibers and blue arrows indicate ectopic divisions in the phloem.

### BP inactivation completely abolishes xylem expansion and secondary growth in *arf6 arf8*

To investigate the interaction and the specific contribution of BP and ARF6 ARF8 during xylem expansion, we generated *arf6 arf8 bp* triple mutants. At 25daf, *arf6 arf8 bp* triple mutant displayed a dramatic reduction of xylem occupancy when compared to *arf6 arf8* double or *bp* mutants (Figures 5a-b), indicating that they act in independent pathways. Moreover, loss of *ARF6* and *ARF8* in *bp* mutants *(arf6 arf8 bp)* leads to a drastic reduction of total hypocotyl area/overall secondary growth compared to *bp* single mutants (Figures 5a-c), suggesting that *ARF6* and *ARF8* regulate cambium activity during all phases of secondary growth. To further clarify this, we investigated cambium development in *arf6 arf8 bp* triple and *arf6 bp* and *arf8 bp* double mutant combinations. At flowering time, both double and triple mutants already displayed reduced xylem area and number of cambial layers when compared to *bp* (Figures 5d-e and S6d), demonstrating that ARF6 and ARF8 promote cambial activity in the absence of BP. Strikingly, cambium development was perturbed to such a degree in *arf6 arf8 bp* mutants that it was not possible to distinguish the typical cambial cell morphology (Figure 5d). Consistently only few secondary xylem vessels were produced (Figure 5f). Altogether these results indicate that ARF6 and ARF8 genetically interact with BP to promote cambium establishment and activity during the early phase of secondary growth.

**Fig. 5.**
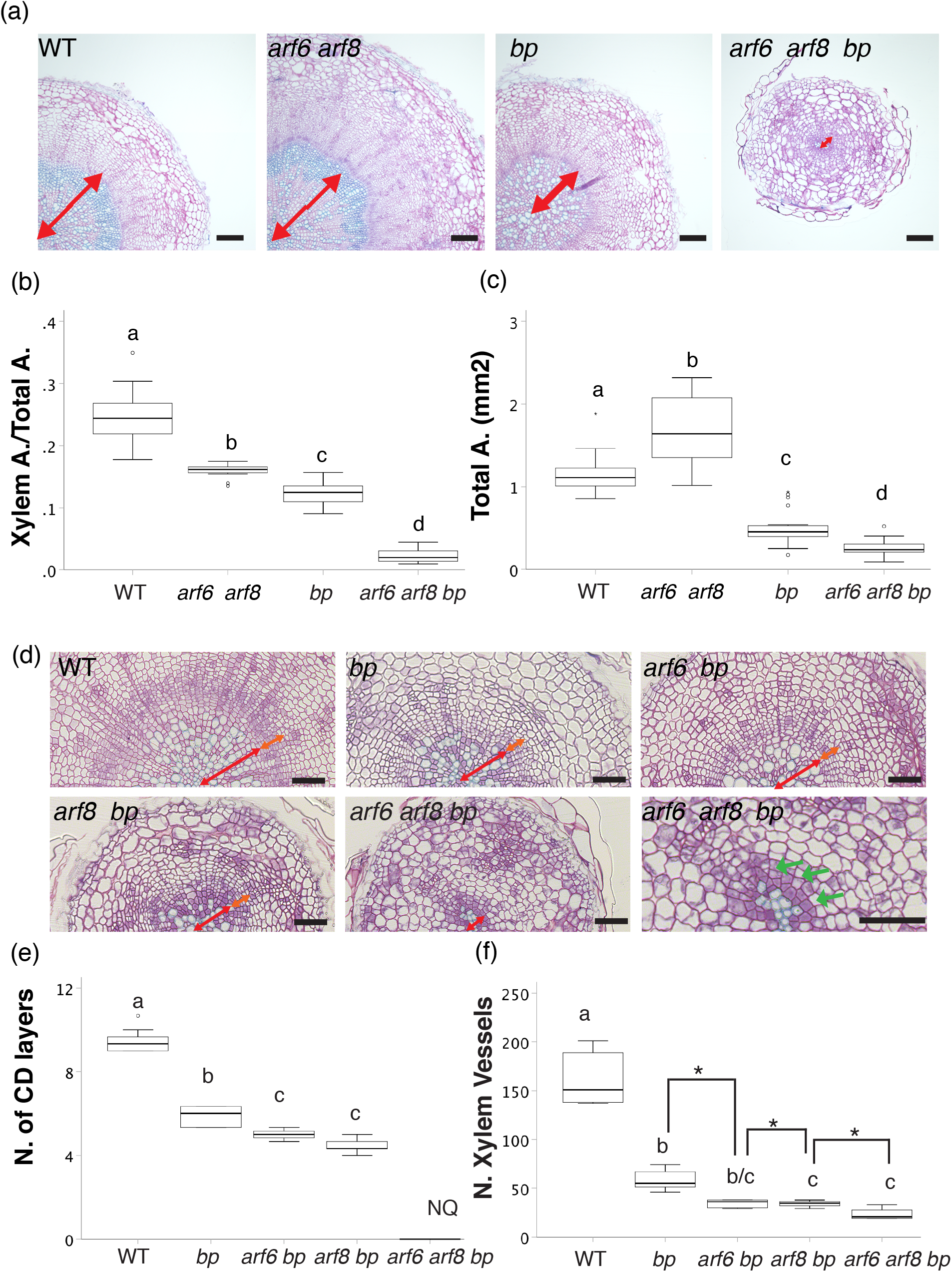
BP inactivation completely abolishes secondary growth in *arf6 arf8*. (a) Plastic hypocotyl cross-sections of 25 day-after-flowering (daf), showing xylem expansion in Col-0, *bp*, *arf6 arf8* and *ar6 arf8 bp* mutants. Black scale bars: 100μm. (b) Quantification of the Xylem Area /Total area ratio of the experiment showed in (a). Letters in the graphs refer to individual groups in a one-way ANOVA analysis with a post-hoc multiple group T-test (n=12-23). (c) Quantification of the Total area of the experiment showed in (a). Letters in the graphs refer to individual groups in a one-way ANOVA analysis with a post-hoc multiple group T-test (n=12-23). (d) Plastic hypocotyl cross-sections of flowering Col-0, *bp*, *arf6 bp, arf8 bp* and *ar6 arf8 bp* mutants. Black scale bars: 50μm. (e) Quantification of the number of cambium derivative layers (CD layers) of the experiment showed in (d). Letters in the graphs refer to individual groups in a one-way ANOVA analysis with a post-hoc multiple group T-test (n=4-7). T-test (* :p<0.05) and NQ (Not possible to quantify). (f) Quantification of the number of xylem vessels of the experiment showed in (d). Letters in the graphs refer to individual groups in a one-way ANOVA analysis with a post-hoc multiple group T-test (n=4-7). Double-headed red arrows indicate the xylem, double-headed orange arrows indicate the cambium and green arrows indicate altered cambial morphology in *arf6 arf8 bp* triple mutant.

## Discussion

Secondary growth results in the thickening of plant organs and formation of secondary xylem (wood), which is a principal sink for carbon and a fundamental source of natural renewable energy (Demura & Ye, 2010). It largely relies on the interdependent processes of cell proliferation and differentiation (Ragni & Greb, 2018). GA positively controls secondary growth by modulating cambial activity and fiber differentiation (Eriksson *et al.*, 2000; Björklund *et al.*, 2007; Ragni *et al.*, 2011; Ikematsu *et al.*, 2017). Differences in GA signaling define two separate phases of secondary growth in Arabidopsis hypocotyl; a first phase in which xylem and phloem are produced at the same rate and a second phase (the so-called xylem expansion) in which xylem production is accelerated and xylem fiber differentiates (Ragni *et al.*, 2011). The transition between these two phases is triggered by local GA-dependent degradation of DELLA proteins, which occurs in the hypocotyl at flowering time (Ragni *et al.*, 2011).

In this study, we further elucidated the GA-mediated DELLA modulation of secondary growth. In Arabidopsis, the *DELLA* gene family comprises 5 members (Zentella *et al.*, 2007) and analyses of *della* single and higher order mutants pointed out that *RGA* and *GAI* are the major contributors. The role of *RGA* and *GAI* in secondary growth is conserved across ecotypes, which greatly differ in xylem expansion dynamics such as Col and Ler. GA signaling, through the degradation of RGA and GAI, triggers xylem expansion and cambium senescence in both accessions and this is independent of *ERECTA*, a known regulator of xylem expansion, which is not functional in Ler background. Interestingly, in both accessions, the expression of undegradable versions of DELLA represses xylem occupancy but only in Col background promotes phloem development and prevents cambium senescence, revealing a new aspect of GA-mediated xylem expansion. This could be explained by the fact that in Ler accession an unknown factor possibly promotes xylem formation independently of DELLA and ERECTA and the resultant high xylem occupancy masks the effect of GA on the cambium /phloem. In line with this idea, a similar scenario has been proposed in the context of flower fertility where an unidentified stamen elongation regulator, which is not ERECTA, promotes stamen elongation independently of DELLA in Ler background (Plackett *et al.*, 2014).

The presence of a xylem promoting factor in Ler accession may also explain why the inactivation of BP in Ler background has a weaker fiber phenotype compared to the Col allele. Despite the major role of BP in promoting fiber formation during xylem expansion, our genetic analyses indicate that BP is not the only DELLA-interacting factor controlling fiber formation. Consistently, DELLA proteins are known to physically interact with many transcription factors belonging to different families (Claeys *et al.*, 2014; Colebrook *et al.*, 2014; Marin-de la Rosa *et al.*, 2014) and act as central hub connecting several signaling cascades such as auxin, cytokinins, ethylene, Jasmonate and ABA in different developmental contexts (Locascio *et al.*, 2013). It is well established that auxin promotes different aspects of secondary growth such as cambium establishment, cambium homeostasis, xylem differentiation and periderm formation (Brackmann *et al.*, 2018; Smetana *et al.*, 2019; Xiao *et al.*, 2020). In this work we show that auxin and GA, via the interaction of ARF6, ARF8 with DELLA, regulate xylem expansion, revealing a novel hormonal cross-talk during secondary growth. ARF6 and ARF8 are known to regulate flower development, adventitious root formation, and hypocotyl elongation (Nagpal *et al.*, 2005; Goetz *et al.*, 2006; Gutierrez *et al.*, 2012; Liu *et al.*, 2014) and recently, we show that ARF8 promotes periderm growth in the root (Xiao *et al.*, 2020). Consistently with their novel role during xylem expansion, we showed that *arf6 arf8* mutants display reduced xylem occupancy, diminished xylary fiber formation, retarded cambium senescence, and enhanced phloem proliferation, mimicking the expression of a undegradable version of DELLA (in Col background). Moreover, GA responses are attenuated in *arf6 arf8* double mutants, and nearly abolished in *arf6 arf7 arf8* triple mutants validating the specific role played by DELLA-ARFs interaction in controlling xylem expansion. In addition*, arf6 arf8* double mutants form phloem fibers, which hardly occurs in WT plants, highlighting the role of ARF6 and ARF8 in repressing phloem development. The presence of phloem fibers is in concordance with recent reports, showing that Jasmonate promotes phloem fiber formation (Behr *et al.*, 2018), as it is known that *ARF6* and *ARF8* activate JA conjugating enzymes, limiting bioactive JA during adventitious root formation (Gutierrez et al 2012). During secondary growth, ARF6 and ARF8 are mainly expressed in the phloem, where also DELLA proteins accumulate, and in the cambium, indicating that the xylem phenotypes of *arf6 arf8* double mutants might be indirect effects of altered cambium/phloem function. These findings highlight, that xylem expansion does not only rely on promoting xylem and xylem fiber formation but also on the repression of phloem proliferation and promoting cambium senescence. To conclude, *ARF6, ARF7, ARF8* and *BP* are sequestered and therefore inactivated by DELLA proteins before flowering, while after flowering (the onset of xylem expansion) bioactive GAs accumulate in the hypocotyl (Talon *et al.*, 1991; Ragni *et al.*, 2011), triggering the degradation of DELLA proteins and the release ARFs and BP (Figure 6). Together ARFs and BP activate the transcriptional programs, which ultimately promotes xylem vessels and fiber production and repression of phloem proliferations.

**Fig. 6.**
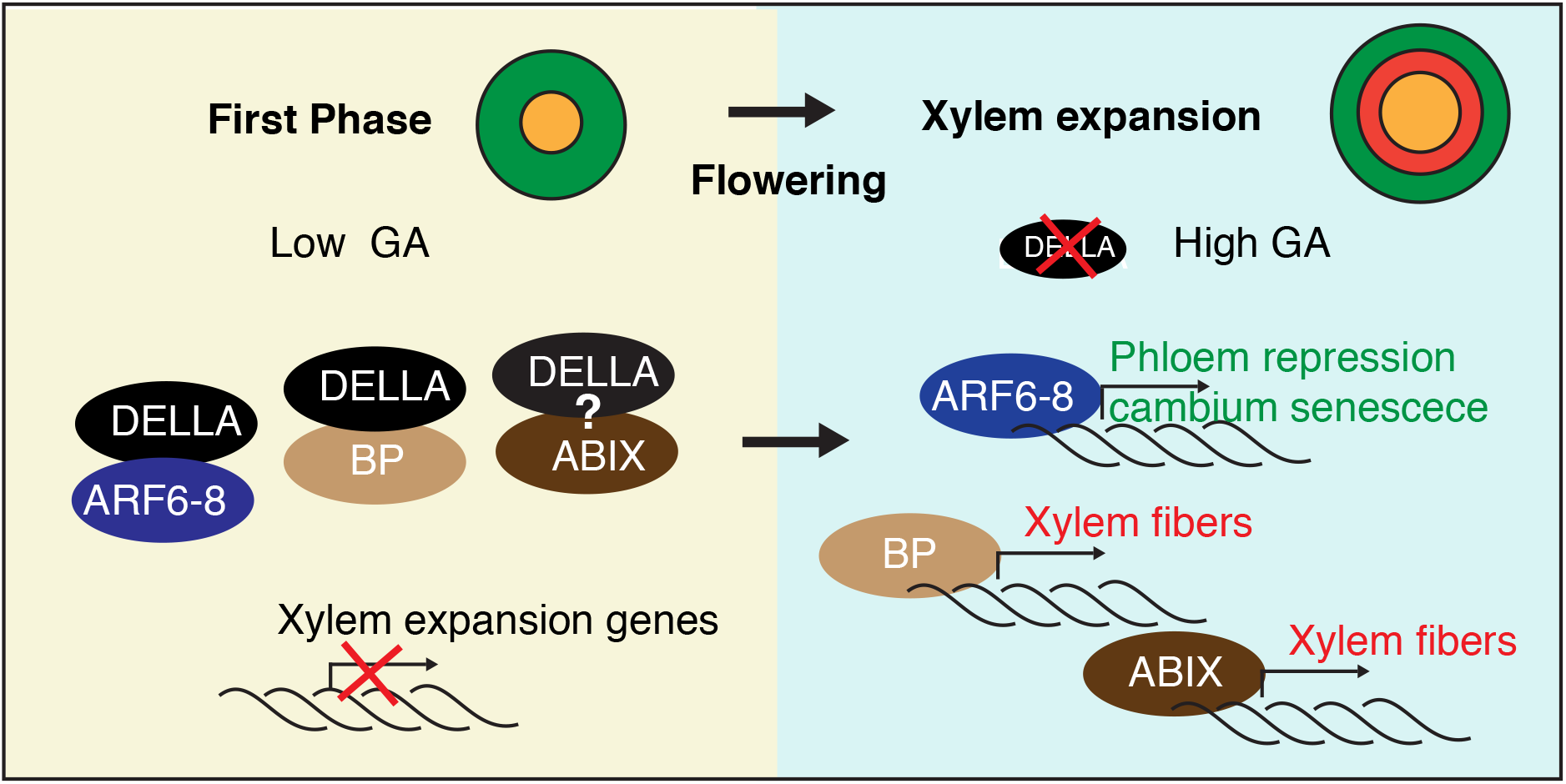
*arf6 arf8* Model explaining GA-mediated xylem expansion. In Arabidopsis hypocotyls, we can distinguish two phases of secondary growth. In the first phase xylem and phloem are produced at the same rate, whereas in the second phase, xylem production is enhanced and xylem fiber differentiated. Before flowering, GA levels are low and DELLA proteins sequester ARFs, BP and probably ABIs, blocking the activation of xylem expansion genes. At flowering, GA accumulates in the hypocotyl and mediates DELLA degradation, freeing ARFs, BP and probably ABIs, which promote cambium senescence, phloem repression and fiber production.

Several ARFs such as ARF5/MONOPTEROS modulate cambium development. In the stem, ARF5 stimulates the transition of cambial stem cells into xylem cells by directly activating xylem-related genes and by repressing the key cambial regulator *WOX4*, whereas ARF3 and ARF4 maintain cambial stem cell proliferation (Brackmann *et al.*, 2018). By contrast, in the root ARF5 with ARF7 and ARF19, promotes the expression of class III HD-ZIPs, *PXY* and *WOX4* triggering the establishment of the vascular cambium (Smetana *et al.*, 2019). Similarly, in the hypocotyl at the onset of secondary growth ARF6 and ARF8 (in the absence of BP) promote cambium activity/ establishment probably by the activation of *WOX4* as *arf6 arf8 bp* triple mutants are reminiscent of *wox4 bp* double mutants (Zhang *et al.*, 2019). Instead, in the late xylem expansion phase ARF6 and ARF8 activate cambial senescence. Consistently, the genetic interaction of *ARF6, ARF8*, and *BP* varies among plant organs, in fact during flower development, *ARF6* and *ARF8* repress *BP* expression and the inactivation of BP partially restores the *arf6 arf8* flower defects (Tabata et al., 2010), whereas in the hypocotyl knocking out BP enhanced the xylem occupancy phenotype of *arf6 arf8.* This is in accordance with other studies showing that BP has different functions in above and underground (root/hypocotyl) organs (Mele et al., 2003; Liebsch et al., 2014; Woerlen et al., 2017).

Interestingly, both BP and ARFs interact with DELLA and are expressed in the phloem highlighting the possibility of higher order protein complex formation. Recently, Campbell et al showed that abscisic acid specifically stimulates fiber formation during xylem expansion independently of BP (Campbell *et al.*, 2018). As DELLAs are known to bind ABA signaling core components such as ABSICS ACID INSENSITIVE 3 and ABI5 (ABI5) (Lim *et al.*, 2013), it is tempting to speculate that the ABA-mediated fiber promotion is regulated by DELLA (Figure 6). Further research focusing on the spatio-temporal dynamics of DELLA, BP, ARFs, and ABI protein accumulation and interaction, coupled with tissue-specific target analyses will help to further dissect the distinct events that lead to xylem expansion, and contrasting functions of ARFs and BP during all stages of secondary growth.

## Supporting information

All supplemental Figures

## Authors contribution

LR, MBT, and MB designed and LR and MBT conducted the experiments, MBT performed the histology with the help of DR. LR, DR and MBT generated the plant lines. LR and MBT wrote the manuscript with the help of MB.

## Acknowledgement

We thank Stefan Mahn for cloning the DELLA promoters and Andrea Boch for helping with plant cultivation. We thank Ari Pekka Mähönen for discussion and reading of the manuscript.

## Supporting Information

**Fig. S1** RGA and GAI are the main DELLA regulating secondary growth.

**Fig. S2** Induction of *rgaD17* and *gaiD17* repress xylem expansion in both Col and Ler ecotypes.

**Fig. S3** Genetic interaction between BP and DELLA.

**Fig. S4** DELLA and ARF expression pattern and *arfs* mutant characterization.

**Fig. S5** Secondary growth dynamics in *arf6 arf8* double mutants.

**Fig. S6** Genetic interaction between ARFs and DELLA.

**Table S1** Primers used for genotyping.

