## Supplementary material for "*AUXIN RESPONSE FACTOR 6* (*ARF6)* and *ARF8* promote Gibberellin-mediated hypocotyl xylem expansion and cambium homeostasis": All supplemental Figures

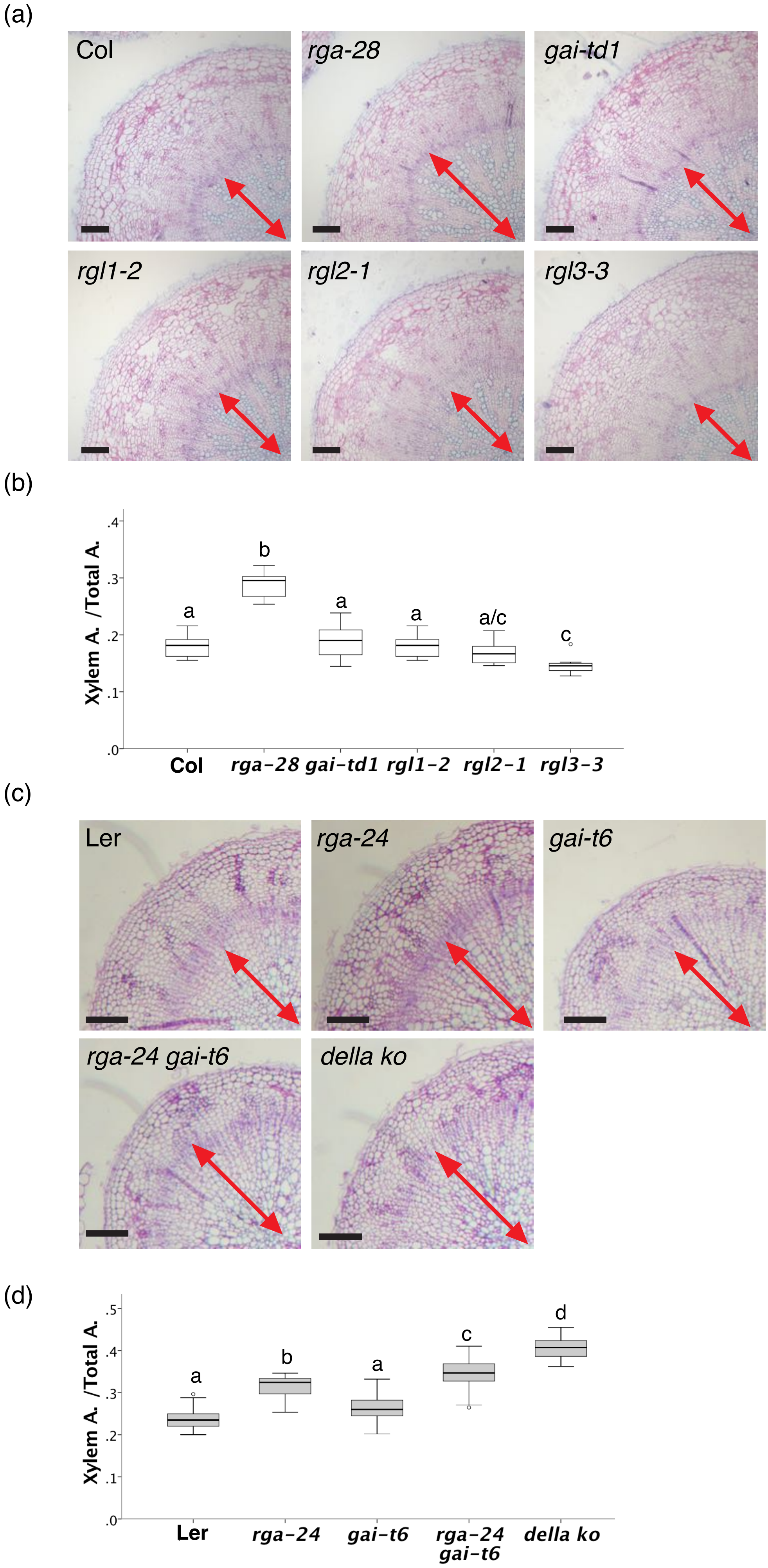

(a)
Col
rga-28
gai-td1
rgl1-2
rgl2-1
rgl3-3
(b)
Xylem A. /Total A.
.4
.3
.2
.1
.0
b
a
a
a
a/c
c
Col
rga-28
gai-td1
rgl1-2
rgl2-1
rgl3-3
(c)
Ler
rga-24
gai-t6
rga-24 gai-t6
della ko
(d)
Xylem A. /Total A.
.5
.4
.3
.2
.1
.0
a
b
a
c
d
Ler
rga-24
gai-t6
rga-24 gai-t6
della ko

**Fig. S1** RGA and GAI are the main DELLA regulating secondary growth.

(a) Plastic hypocotyl cross-sections of 10 day-after-flowering (daf), showing xylem expansion in della single mutants (in Col background): *rga-28*, *gai-td1*, *rgl1-2*, *rgl2-1* and *rgl3-3*. (b) Quantification of the Xylem Area /Total area ratio of the experiment showed in (a). Letters in the graphs refer to individual groups in a one-way ANOVA analysis with a post-hoc multiple group T-test (n=11-12). (c) Plastic cross-sections of 8daf, showing xylem expansion in different combination of della mutants (in Ler background): *rga-24*, *gai-t6*, *rga-24 gai-t6* and *della ko*. (d) Quantification of the Xylem Area /Total area ratio of the experiment showed in (c). Letters in the graphs refer to individual groups in a one-way ANOVA analysis with a post-hoc multiple group T-test (n=20) (c). Black scale bar: 100µm. Double-headed red arrows indicate the xylem.

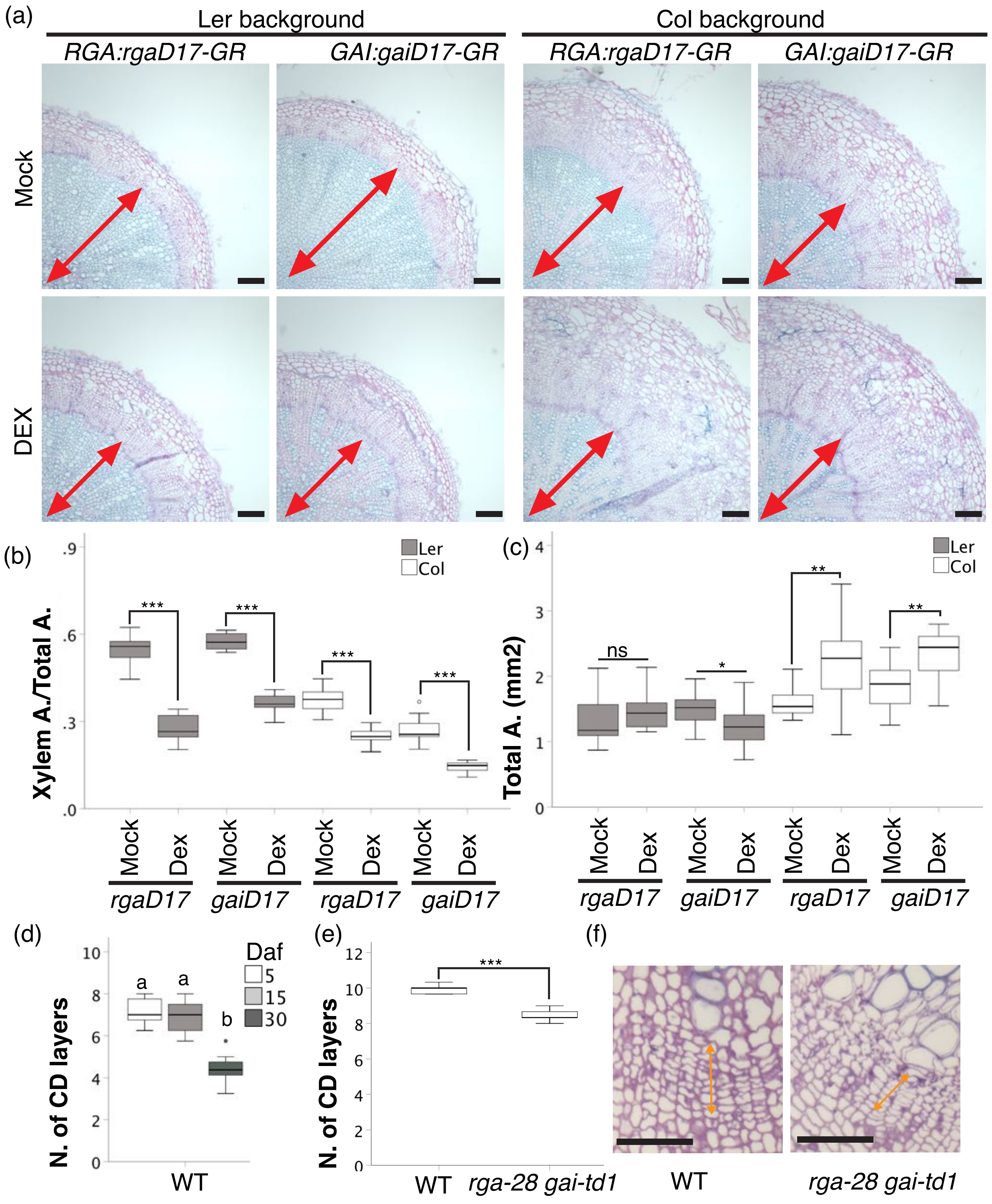

(a)
Ler background
Col background
RGA:rgaD17-GR
GAI:gaiD17-GR
RGA:rgaD17-GR
GAI:gaiD17-GR
Mock
DEX
(b)
Xylem A./Total A.
Ler
Col
***
Mock Dex Mock Dex Mock Dex Mock Dex
rgaD17 gaiD17 rgaD17 gaiD17
(c)
Total A. (mm2)
ns
*
**
Mock Dex Mock Dex Mock Dex Mock Dex
rgaD17 gaiD17 rgaD17 gaiD17
(d)
N. of CD layers
Daf
5
15
30
a a b
WT
(e)
N. of CD layers
***
WT rga-28 gai-td1
(f)
WT
rga-28 gai-td1

**Fig S2** Induction of *rgaD17* and *gaiD17* repress xylem expansion in both Col and Ler ecotypes. (a) Plastic cross-sections of 20 day-after-flowering (daf) hypocotyls of *rga:rgaD17-GR* and *gai:gaiD17-GR* in Col and Ler backgrounds. Plants were treated with 10µM DEX from flowering until sample collection. Double-headed red arrows indicate the xylem. Black scale bars:100µm. (b) Quantification of Xylem Area/Total area of the experiment showed in (a). T-test (n=11-13, ***: $p<0.001$). (c) Quantification of Total Area of the experiment showed in (a). T-test (n=11-13, ns: Not significant, * : $p<0.05$, **: $p<0.01$). (d) Quantification of the number of cambium derivative layers (CD layers) in the hypocotyl of WT plants at 5, 15, 30 daf. Letters in the graphs refer to individual groups in a one-way ANOVA analysis with a post-hoc multiple group T-test (n=6). This experiment was also used as WT control for Figure S5e. (e) Quantification of the number of cambium derivative layers (CD layers) of the experiment showed in (f). T-test (n=6, ***$<0.001$). (f) Magnifications of WT and *rga-28- gai-td1* hypocotyl crosse-sections relative to Figure 1a. Double-headed orange arrows indicate the cambium. Black scale bar: 20µm.

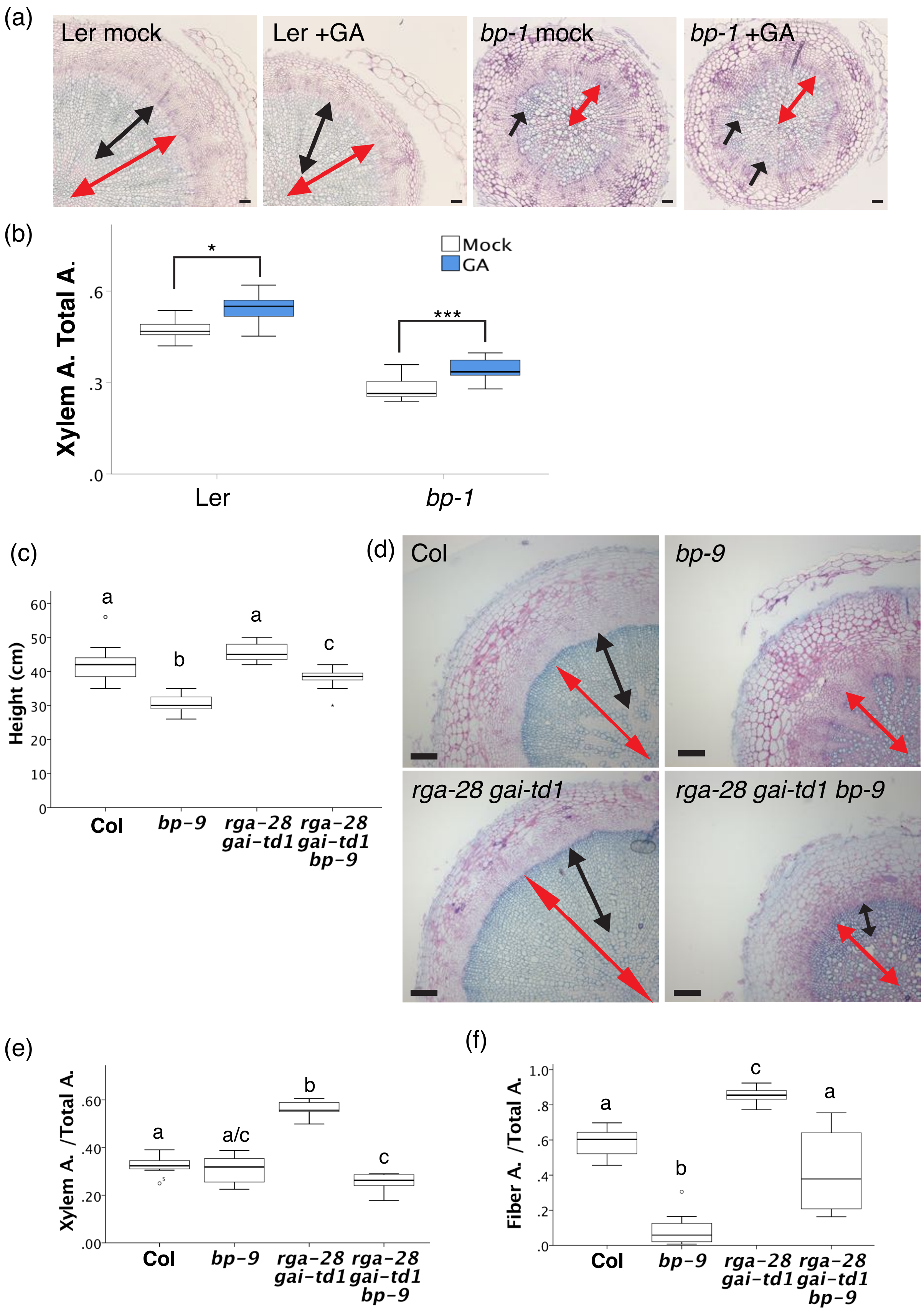
(a)
Ler mock
Ler +GA
bp-1 mock
bp-1 +GA
(b)
Mock
GA
Xylem A. Total A.
.6
.3
.0
*
***
Ler
bp-1
(c)
Height (cm)
60
50
40
30
20
10
0
a
b
a
c
Col
bp-9
rga-28 gai-td1
rga-28 gai-td1 bp-9
(d)
Col
bp-9
rga-28 gai-td1
rga-28 gai-td1 bp-9
(e)
Xylem A. /Total A.
.60
.40
.20
.00
a
a/c
b
c
Col
bp-9
rga-28 gai-td1
rga-28 gai-td1 bp-9
(f)
Fiber A. /Total A.
1.0
.8
.6
.4
.2
.0
a
b
c
a
Col
bp-9
rga-28 gai-td1
rga-28 gai-td1 bp-9

**Fig. S3** Genetic interaction between BP and DELLAs.

(a) Plastic cross-sections of 20 day-after-flowering (daf) hypocotyls of *Ler* and *bp-1*. Plants were treated with 10µM GA from flowering until sample collections. Black scale bars:100µm. (b) Quantification of Xylem Area/Total area of the experiment showed in (a). T-test (n=18-15, *: $p<0.05$, *** $p<0.001$). (c) Quantification of stem height in WT (Col), *bp-9*, *rga-28*, *gai-td1*, *rga-28 gai-td1 bp-9* at plant senescence. Letters in the graphs refer to individual groups in a one-way ANOVA analysis with a post-hoc multiple group T-test (n=20). (d) Plastic cross-sections of 25daf hypocotyls of Col, *bp-9*, *rga-28*, *gai-td1*, *rga-28 gai-td1 bp-9*. (e) Quantification of Xylem Area/Total area of the experiment showed in (d). Letters in the graphs refer to individual groups in a one-way ANOVA analysis with a post-hoc multiple group T-test (n=8-9). (f) Quantification of Fiber Area/Xylem area of the experiment showed in (d). Letters in the graphs refer to individual groups in a one-way ANOVA analysis with a post-hoc multiple group T-test (n=8-9). Double-headed red arrows indicate the xylem, double-headed and single-headed black arrows indicate xylem fibers. Black scale bars:100µm.

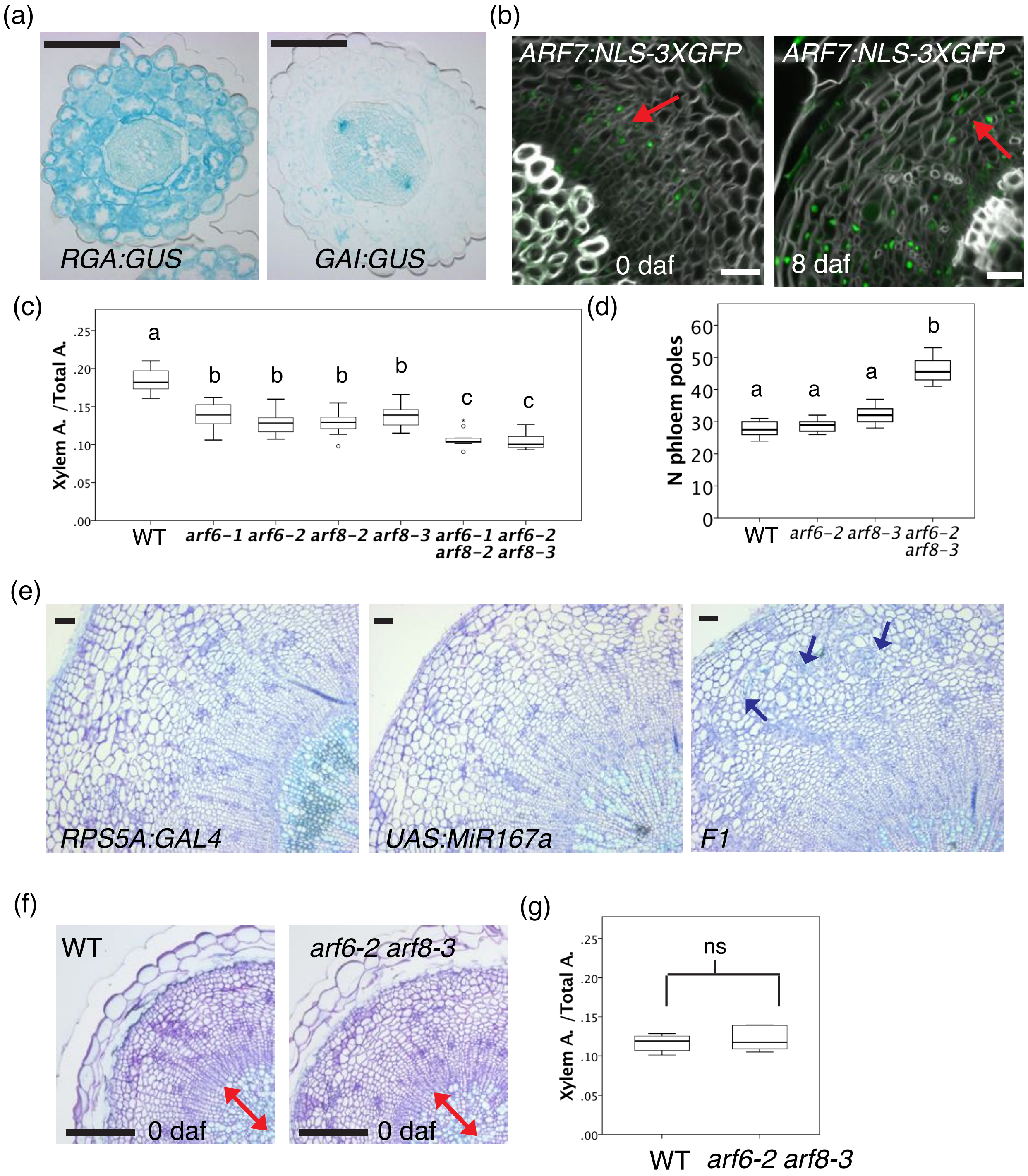

(a)
RGA:GUS
GAI:GUS
(b)
ARF7:NLS-3XGFP
0 daf
ARF7:NLS-3XGFP
8 daf
(c)
Xylem A. /Total A.
.25
.20
.15
.10
.05
.00
a
b
b
b
b
c
c
WT
arf6−1
arf6−2
arf8−2
arf8−3
arf6−1 arf8−2
arf6−2 arf8−3
(d)
N phloem poles
60
50
40
30
20
10
0
a
a
a
b
WT
arf6−2
arf8−3
arf6−2 arf8−3
(e)
RPS5A:GAL4
UAS:MiR167a
F1
(f)
WT
0 daf
arf6-2 arf8-3
0 daf
(g)
Xylem A. /Total A.
.25
.20
.15
.10
.05
.00
ns
WT
arf6-2 arf8-3

**Fig. S4** DELLA and ARF expression pattern and *arfs* mutant characterization.

(a) Plastic hypocotyl cross-sections of *RGA:GUS* and *GAI:GUS* at 16 days after germination. (before flowering and the onset of xylem expansion). (b) Vibratome hypocotyl cross-sections of *ARF7:NLS-GFP-GUS* respectively at 0 and 8 day-after-flowering (daf). Red arrows indicate GFP signal in the phloem. (c) Quantification of Xylem/Total area ratio in 10 daf hypocotyls of WT (Col), *arf6-1*, *arf6-2*, *arf8-2*, *arf8-3*, *arf6-1 arf8-2* and *arf6-2 arf8-3*. Letters in the graphs refer to individual groups in a one-way ANOVA analysis with a post-hoc multiple group T-test (n=8-10). (d) Quantification of the number of phloem poles in WT (Col), *arf6*, *arf8* and *arf6 arf8* at 10 daf relative to Figure 3a. Letters in the graphs refer to individual groups in a one-way ANOVA analysis with a post-hoc multiple group T-test (n=6) (e) Plastic hypocotyl cross-sections of 10 daf *RPS5A:GAL4*, *UAS:MIR167a* and F1 (*RPS5A:GAL4 x UAS:MIR167a*). Blue arrows indicate ectopic divisions in the phloem. (f) Plastic hypocotyl cross-sections of flowering plants showing xylem expansion in WT (Col) and *arf6-2 arf8-3* double mutants. Double-headed red arrows indicate the xylem. (g) Quantification of the Xylem Area /Total area ratio of the experiment showed in (f). T-test (n=6-8, ns: not significant). White scale bars: 20 µm, black scale bar:100µm.

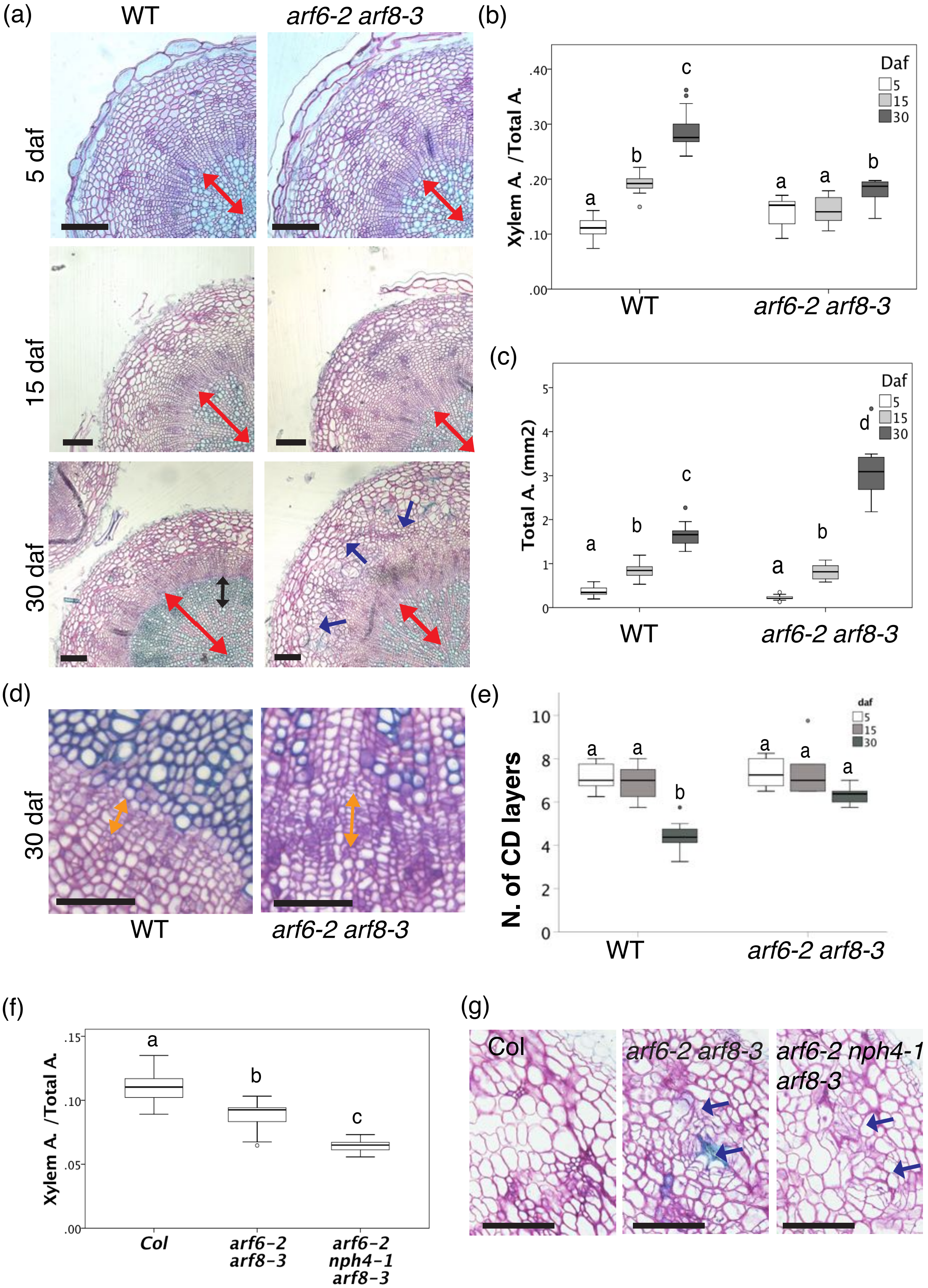

(a)
WT
arf6-2 arf8-3
5 daf
15 daf
30 daf
(b)
Xylem A. /Total A.
Daf
5
15
30
WT
arf6-2 arf8-3
(c)
Total A. (mm2)
Daf
5
15
30
WT
arf6-2 arf8-3
(d)
30 daf
WT
arf6-2 arf8-3
(e)
N. of CD layers
daf
5
15
30
WT
arf6-2 arf8-3
(f)
Xylem A. /Total A.
Col
arf6-2 arf8-3
arf6-2 nph4-1 arf8-3
(g)
Col
arf6-2 arf8-3
arf6-2 nph4-1 arf8-3

**Fig. S5** Secondary growth dynamics in *arf6 arf8* double mutants.

(a) Plastic hypocotyl cross-sections showing xylem expansion in WT (Col) and *arf6-2 arf8-3* double mutants at 5, 15, 30 day-after- flowering (daf). (b) Quantification of the Xylem Area /Total area ratio of the experiment showed in (a). Letters in the graphs refer to individual groups in a one-way ANOVA analysis with a post-hoc multiple group T-test (n=10-14). (c) Quantification of the Total area in the experiment of the experiment showed in (a). Letters in the graphs refer to individual groups in a one-way ANOVA analysis with a post-hoc multiple group T-test (n=10-14). (d) Magnifications showing the cambium of WT (Col) and *arf6-2 arf8-3* double mutants at 30 daf relative to Figure S5a. (e) Quantification of the number of cambium derivative layers (CD layers) of the experiment showed in (d). The same experiment (for the WT) is presented in Figure S2d. Letters in the graphs refer to individual groups in a one-way ANOVA analysis with a post-hoc multiple group T-test (n=6). (f) Quantification of the Xylem Area /Total area ratio of the experiment showed in Figure 3f. Letters in the graphs refer to individual groups in a one-way ANOVA analysis with a post-hoc multiple group T-test (n=8-11). (g) Magnifications of cross-sections showing the phloem of WT (Col), *arf6-2 arf8-3* and *arf6-2 nph4-1 arf8-3* at 10 daf relative to figure 3f. Double-headed red arrows indicate the xylem, double-headed orange arrows indicate the cambium, double-headed black arrows indicate xylem fibers and blue arrows indicate ectopic divisions in the phloem. Black scale bars:100µm.

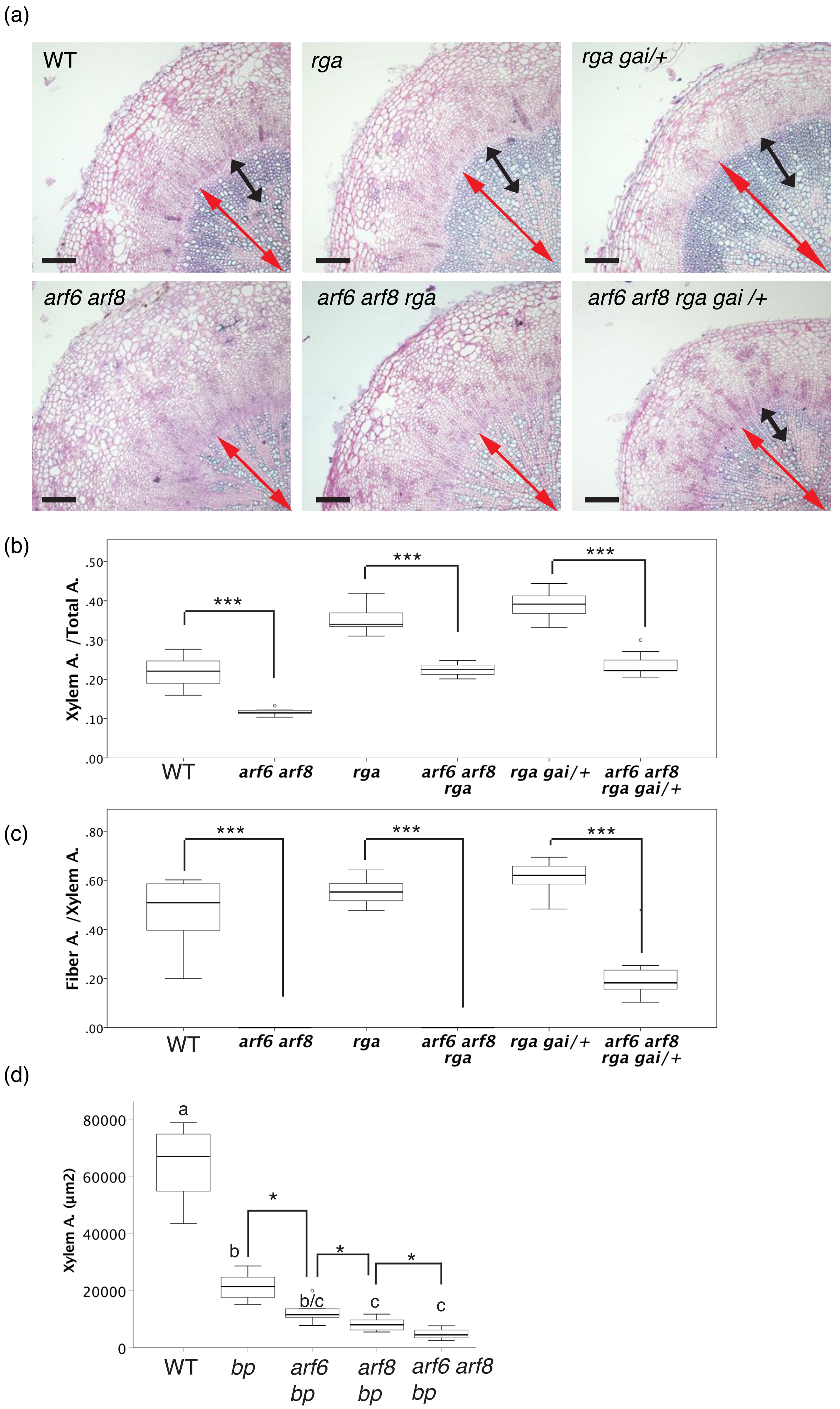

(a)
WT
rga
rga gai/+
arf6 arf8
arf6 arf8 rga
arf6 arf8 rga gai /+
(b)
Xylem A. /Total A.
.50
.40
.30
.20
.10
.00
***
***
***
WT
arf6 arf8
rga
arf6 arf8 rga
rga gai/+
arf6 arf8 rga gai/+
(c)
Fiber A. /Xylem A.
.80
.60
.40
.20
.00
***
***
***
WT
arf6 arf8
rga
arf6 arf8 rga
rga gai/+
arf6 arf8 rga gai/+
(d)
Xylem A. (µm2)
80000
60000
40000
20000
0
a
b
b/c
c
c
*
*
*
WT
bp
arf6 bp
arf8 bp
arf6 arf8 bp

**Fig. S6** Genetic interaction between ARFs and DELLA.

a) Plastic hypocotyl cross-sections stained, showing xylem expansion in WT (Col), *arf6 arf8*, *rga gai/+*, *arf6 arf8 rga* and *arf6 arf8 rga gai/+* at 15 day-after-flowering (daf).(b) Quantification of Xylem/Total area ratio of the experiment showed in (a). T-test (n=3,12, ***: P < 0.001). (c) Quantification of Fiber area/Xylem area ratio of the experiment showed in (a). T-test (n=3-12, ***: P < 0.001). (d) Quantification of the Xylem Area of the experiment showed in Figure 5d. Letters in the graphs refer to individual groups in a one-way ANOVA analysis with a post-hoc multiple group T-test (n= 4-6). T-test (n= 4-6), *: P < 0.05). Double-headed red arrows indicate the xylem and double-headed black arrows indicate xylem fibers. Black scale bars:100µm.

**Table S1** Primers used for genotyping.

| Allele | WT/mut | primer name | sequence |
|---|---|---|---|
| *rga-28* | WT | RGA genoF | CGATTGTCCAACCACGGG |
| | | RGAR_201 | CAGCTAAGCATCCGATTTGC |
| | mut | rga28-244 | ATGGCGGAGGTTGCTTTGAAACTCGAACA |
| | | Ds3-2 | CCGGTATATCCCGTTT TCG |
| *gai-td1* | WT | GAI_TDNA_LP3 | CGGTAACGGCATGGATGAG |
| | | GAI_TDNA_RP | AGCTTCGGCGAAGTAAGTAGC |
| | mut | GAI_TDNA_RP | AGCTTCGGCGAAGTAAGTAGC |
| | | LB3 | TAGCATCTGAATTTCATAACCAATCTCGATACAC |
| *rgl1-1* | WT | RGL1 1670F | AAGCTAGCTCGAAACCCCAAAT |
| | | RGL1 2295R | CCACAGAGCGCGTAGAGGATAAC |
| | mut | RGL1 1670F | AAGCTAGCTCGAAACCCCAAAT |
| | | DS5-P1 | CATGGGCTGGGCCTCAGTG |
| *rgl2-1* | WT | RGL2 856F | GCTGGTGAAACGCGTGGGAACA |
| | | RGL2 1883R | ACGCCGAGGTTGTGATGAGTG |
| | mut | RGL2 856F | GCTGGTGAAACGCGTGGGAACA |
| | | DS5-3 | CGGTCGGTACGGGATTTTCC |
| *arf6-1* | WT | arf6-2 R | CCAAGGGTCATCGCCGAGGAGAAGAACGTC |
| | | arf6-2 F | GACGAATCTACTGCAGGAG |
| | mut | arf6-2 F | GACGAATCTACTGCAGGAG |
| | | LBb1.3 | ATTTTGCCGATTTCGGAAC |
| *arf6-2* | WT | arf6-2 R | CCAAGGGTCATCGCCGAGGAGAAGAACGTC |
| | | arf6-2 F | GACGAATCTACTGCAGGAG |
| | mut | arf6-2 F | GACGAATCTACTGCAGGAG |
| | | JMLB | GGCAATCAGCTGTTGCCCGTCTCACTGGTG |
| *arf8-2* | WT | arf8-3 R | CCATGGGTCATCACCAAGGAGAAGAATATC |
| | | arf8-7 F | CAGGGCTAGCCAATCTGAGTTTGTGATACA |
| | mut | arf8-7 F | CAGGGCTAGCCAATCTGAGTTTGTGATACA |
| | | LB3 | TAGCATCTGAATTTCATAACCAATCTCGATACAC |
| *arf8-3* | WT | arf8-7 F | CAGGGCTAGCCAATCTGAGTTTGTGATACA |
| | | arf8-7 R | GACCACTTCCCAAATCACCCCTTCCATCTG |
| | mut | arf8-7 R | GACCACTTCCCAAATCACCCCTTCCATCTG |
| | | JMLB | GGCAATCAGCTGTTGCCCGTCTCACTGGTG |
| *arf8-7* | WT | ARF8-7genoF | TAAACTTCCATTCAACATCATGGA |
| | | ARF8-7genoR | AGTCGAGTTGTTTACTTTCCACAG |
| | mut | ARF8-7genoR | AGTCGAGTTGTTTACTTTCCACAG |
| | | 8474 | ATAATAACGCTGCGGACATCTACATTTT |
| *nph4-1* | WT | nph4-1geno_F | TCCTGCTGAGTTTGTGGTTCCTT |
| | | nph4-1geno_R | GGGGCTTGCTGATTCTGTTTGTTA |
| | mut | nph4-1geno_R | GGGGCTTGCTGATTCTGTTTGTTA |
| | | LBb1.3 | ATTTTGCCGATTTCGGAAC |
| *bp-9* | WT | BP-14 | TGTTAAGGGTTAGAACACCATG |
| | | BP-3 | GACAACAGCACCACTCCTCAAA |
| | mut | BP-3 | GACAACAGCACCACTCCTCAAA |
| | | dspm1 | CTTATTTCAGTAAGAGTGTGGGGTTTTGG |
